# Optimizing the conservation of migratory species over their full annual cycle

**DOI:** 10.1101/268805

**Authors:** R. Schuster, S. Wilson, A.D. Rodewal, P. Arcese, D. Fink, T. Auer, J.R Bennett

## Abstract

Limited knowledge of the distribution, abundance, and habitat associations of migratory species introduces uncertainty about the most effective conservation actions. We used Neotropical migratory birds as a model group to evaluate contrasting approaches to land prioritization to support ≥30% of the global abundances of 117 species throughout the annual cycle in the Western hemisphere. Conservation targets were achieved in 43% less land area in plans based on annual vs. weekly optimizations. Plans agnostic to population structure required comparatively less land area to meet targets, but at the expense of representation. Less land area was also needed to meet conservation targets when human-dominated lands were included rather than excluded from solutions. Our results point to key trade-offs between efforts minimizing the opportunity costs of conservation vs. those ensuring spatiotemporal representation of populations, and demonstrate a novel approach to the conservation of migratory species based on leading-edge abundance models and linear programming to identify portfolios of priority landscapes and inform conservation planners.

---

Land-use change is a key threat to the conservation of biodiversity, ecosystems^1^, and the services they provide globally^2,3^, and migratory species are particularly vulnerable to such change given the vast geographic areas they occupy over the annual cycle^4,5^. Indeed, a recent global assessment indicated that protected areas adequately protect the ranges of just 9% of migratory bird species^5^. Strategic approaches to identify and conserve habitats critical to the persistence of migratory species are therefore sorely needed.

Unfortunately, substantial gaps in knowledge of the abundance, distribution, and demography of most migratory species^6^ have hampered strategic planning and led to uncertainty about the optimal allocation of conservation effort^5,7^. Given that populations of many migratory species continue to decline^4,8^, there is an urgent need to identify portfolios of lands critical to the persistence of target species, and amenable to management in support of species conservation without compromising human well-being.

Multi-species decision support tools can facilitate the identification of areas crucial to the conservation of migratory species but have remained intractable due to limits on knowledge and computing power. We capitalized on advances in crowd-sourced models of bird species abundance and distribution^9,10^ and linear programing techniques^11^ to develop a robust multi-species planning tool to estimate the land area needed to conserve 117 Nearctic-Neotropical migratory songbirds throughout the annual cycle (SI Table 1). Specifically, we combined fine-scale, predictive models of distribution and abundance estimated weekly throughout the year with spatial optimization techniques^12^ to identify the amount and type of land needed to reach our conservation targets given alternative planning scenarios at hemispheric scales.

**Table 1.**
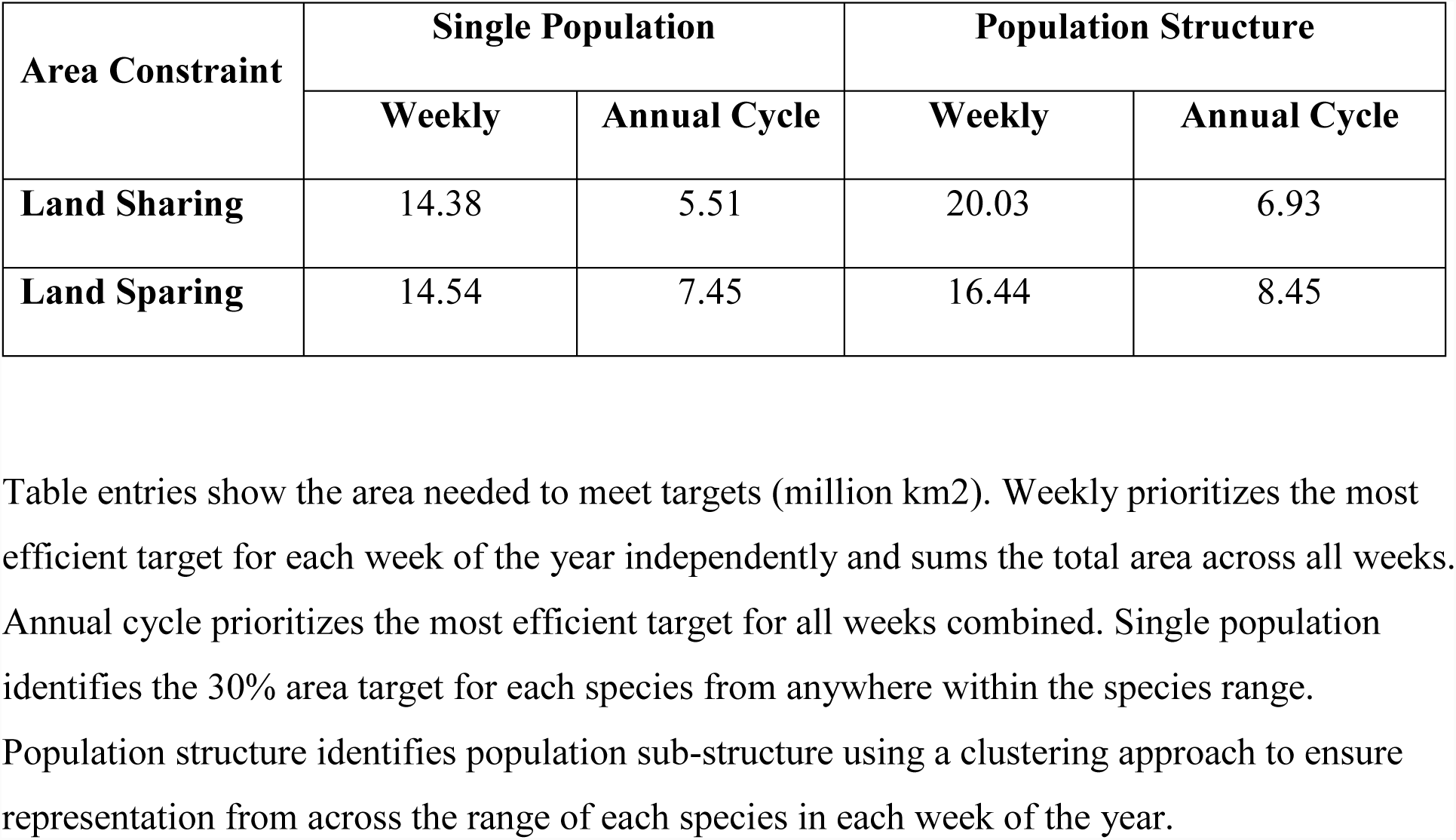
Area requirements to meet a 30% population target for 117 Neotropical migrant bird species for different prioritization approaches under land sharing and land sparing scenarios.

We first estimated the abundance and distribution of 117 migratory bird species weekly, using spatiotemporal exploratory models^9,13^ to calculate the relative abundance of each species throughout the annual cycle (SI Fig. 1). We next recorded and compared the geographic area requirements and land cover types selected when optimizing during each week of the annual cycle (hereafter, “weekly”), versus simultaneously over the entire annual cycle (hereafter, “full annual cycle”). Because all existing conservation plans consider stationary phases of the breeding and non-breeding periods separately^14,15^, our analysis is the first example of spatial optimization scenarios which track populations over their full annual cycle.

**Figure 1.**
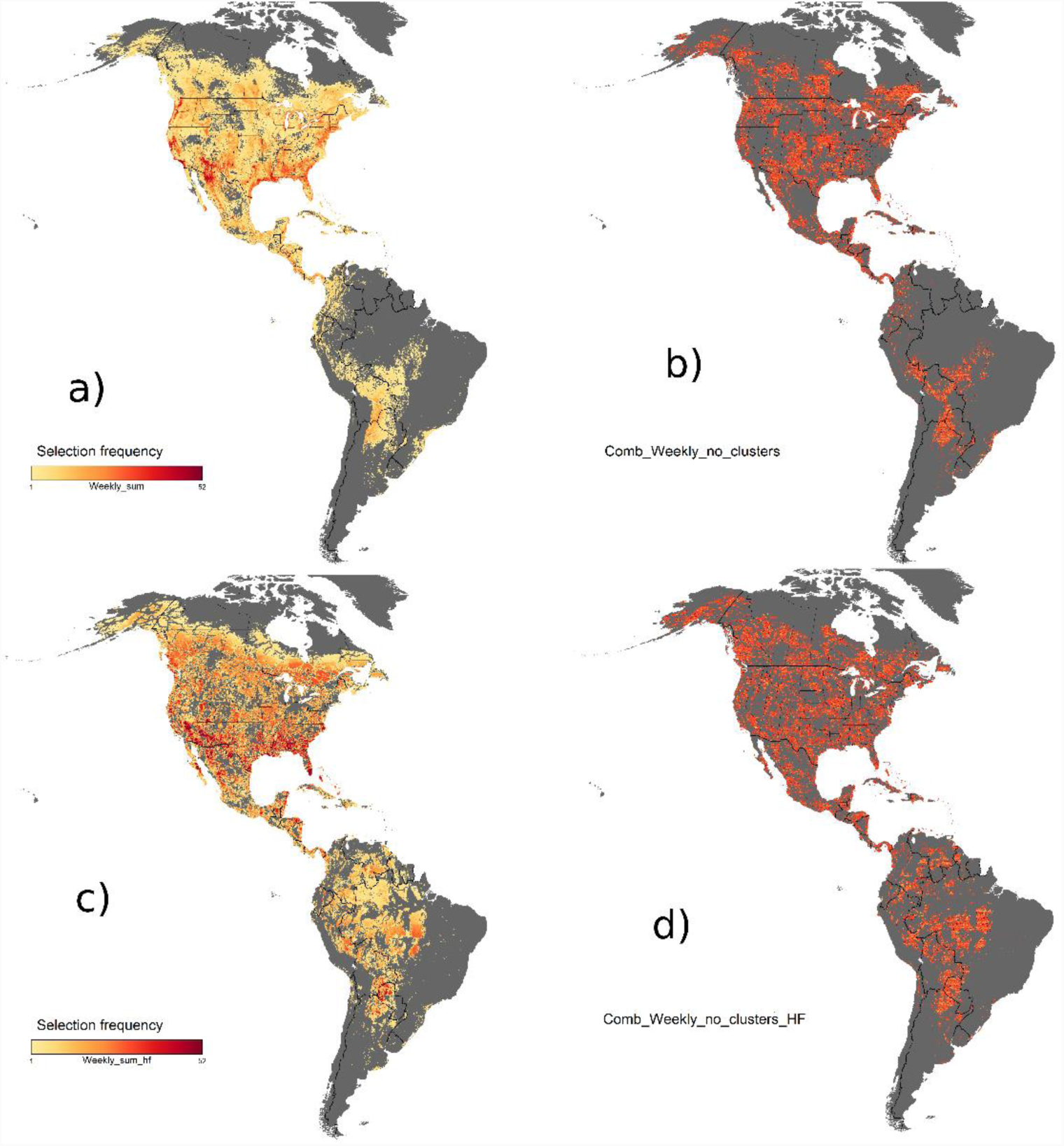
Comparison of areas prioritized for weekly and full annual cycle planning under a land sharing approach allowing for the inclusion of human dominated landscapes versus a land sparing approach that excludes areas of high human footprint. The prioritization is based on a target of 30% of global populations of 117 species of Neotropical migratory birds when each species range is considered as a single population. a) = land sharing, weekly, b) = sharing annual cycle, c) = land sparing weekly, d) = land sparing annual cycle. A more detailed version of this figure focusing on northern South America is SI Figure 4.

We next created area-optimized solutions designed to conserve lands used by 30% of the global populations of all 117 species in each of 52 weeks by sampling species a) over their entire range, without accounting for population structure, or b) by sampling within 5 regional population clusters, identified weekly to accommodate spatial variation in population structure and migratory connectivity. Our 30% target is arbitrary, but intermediate to the 17% of terrestrial ecosystems targeted by the Convention on Biodiversity^16^ and 50% targets suggested by comparative analysis^17^, and it can be easily modified to reflect strategic goals^18^.

Last, we compared area-based conservation plans designed to represent different perspectives about the potential contribution of human-modified lands to the conservation of migratory birds. Our ‘land-sparing’ approach emphasized the protection of relatively intact habitat as indicated by a low human footprint index^19^ (SI Fig. 2), whereas our ‘land sharing’ approach permitted the inclusion of landscapes converted to more intensive use by humans^20^. Exploring such constraints represents a critical step in conservation planning, given that human cultural history, values, and well-being can all affect conservation success and represent critical inputs into structured decisions about the most efficacious actions^21–23^.

**Figure 2.**
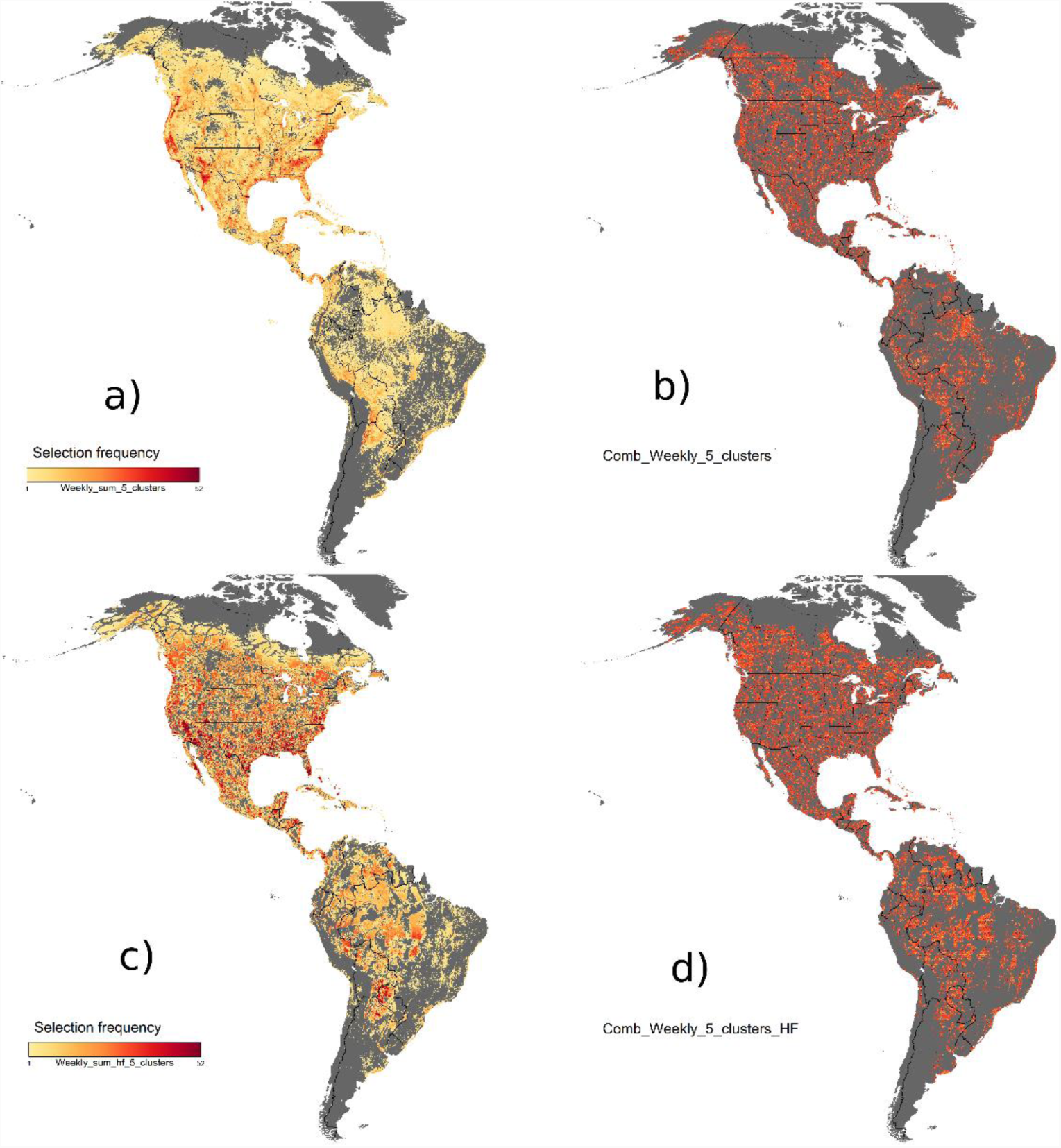
Comparison of areas prioritized for weekly and full annual cycle planning under a land sharing approach allowing for the inclusion of human dominated landscapes versus a land sparing approach that excludes areas of high human footprint. The prioritization is based on a target of 30% of global populations of 117 species of Neotropical migratory birds when each species range is considered with population structure (five regional clusters). a) = land sharing, weekly, b) = sharing annual cycle, c) = land sparing weekly, d) = land sparing annual cycle. A more detailed version of this figure focusing on northern South America is SI Figure 5.

## Results and Discussion

The land area required to achieve conservation targets declined by 56% on average when prioritizations were conducted over the full annual cycle rather than weekly (range = 49% to 65%; Table 1). Full annual cycle solutions also resulted in less land area being prioritized in land-sharing and land-sparing scenarios as compared to solution based on weekly approaches (62% and 49%, respectively; Table 1, Fig. 1, 2). These area reductions under full annual cycle planning generally result from cases such as the inclusion of sites used by a single species in two or more weeks of the year, or by two or more species in during two or more weeks.

Because population structure – let alone its consequences for movement or connectivity – is poorly understood in most migratory species^24^, we developed an innovative approach to account for structure statistically. Specifically, we delineated the populations of each species into 5 spatial clusters and stratified our weekly sampling among clusters to capture the full geographic distribution of each species. As expected, the area required to reach our conservation targets increased when we accommodated population structure, though relatively less so under a land-sparing (13% increase) compared to a land-sharing (26% increase) scenario (Table 1, Figs. 1, 2). Although we currently lack empirical data with which our spatial clusters can be validated, our predictions can be tested directly as tracking and genetic mapping techniques improve to allow comparisons of observed and predicted migration routes. That said, our current method provides a useful approach to ensure geographic representation of population structure of a broad suite of species using publicly-available citizen science data in spatial planning tools.

Many conservation interventions, including land protection, are constrained by limits on fiscal or human resources and the opportunity costs of development. Our results indicate that land area represents one of the key trade-offs in conservation designed to account for population structure and migratory connectivity. In particular, we showed that sampling populations across the species range each week required almost twice the amount of land compared to plans based on the relative abundance of species mapped over the full annual cycle. Our work thus offers the first empirical evidence to support recent calls to assess conservation needs of migratory species across the annual cycle in ways that conserve regional representation, species diversity, and adaptive potential^5,7,10,25^. These findings suggest a need to re-evaluate conservation planning processes based on less precise methods. For example, government and non-governmental organizations allocate up to $1 billion annually to bird conservation based on aspatial targets and expert elicitation, with most actions directed to breeding habitat^14,15^. Our results suggest an alternative approach that stands to meet conservation targets at lower land management cost and, potentially, more compatible with human-dominated lands potentially serving a dual purpose of supporting migratory species and human livelihoods.

Another key result of our work is that incorporating conservation objectives in human-dominated habitats may dramatically improve the efficiency of conservation area designs if the demographic performance of migrants is similar in ‘working’ and ‘intact’ landscapes. We found that land-sharing approaches required 27% and 18% less land area, respectively, than land-sparing approaches including or ignoring population structure (Table 1). Our findings thus add to a growing body of literature indicating the need to broaden the lens through which we view conservation to both accommodate human livelihoods and conserve valued species^21–23^.

Our comparisons of land-sharing and land-sparing approaches identified other geographical or ecosystem-related factors that might influence conservation decisions. Most notably, land-sparing approaches selected larger areas of needle-leaved forest in boreal and mountainous zones of western Canada, and more broad-leaved evergreen forest in the eastern Andes and western Amazon basin (Fig. 1,2; Table 2). Weekly and full annual cycle approaches to land-sparing resulted in geographically similar outcomes (Fig. 1,2), but also differed in land cover types selected (Table 2). Whereas annual cycle planning with land sparing consistently increased the amount of land prioritized over most types of land cover, weekly approaches with and without land sparing resulted in large increases in area requirement for some cover-types and decreases in others (e.g., Table 2 and Table S1). In particular, a weekly, land-sparing approach favored broadleaf evergreen over mixed and broadleaf deciduous forest. Overall, these results illustrate potential trade-offs that conservation practitioners considering optimized portfolios must consider as additional targets and constraints are identified and incorporated in higher-level management models^23^. Even without consensus among conservation practitioners on which scenario to focus on, there is still a considerable amount of land selected in at least six of the eight scenarios investigated, illustrating priority areas that most approaches agree on (126, 000 km^2^, Figure 3).

**Table 2.** Area selected (1000 km^2^) for major land cover types using full annual cycle planning for land sharing vs. sparing scenarios and for single population vs population structure approaches.

| Land cover | Area available | Single Population |  |  | Population Structure |  |  |
| --- | --- | --- | --- | --- | --- | --- | --- |
|  |  | Land Sparing | Land Sharing | % reduction | Land Sparing | Land Sharing | % reduction |
| Cropland/Mosaic Cropland | 2269 | 339 | 313 | 8 | 445 | 439 | 1 |
| Grassland | 5555 | 1198 | 1088 | 9 | 1313 | 1238 | 6 |
| Urban areas | 205 | 9 | 95 | -956 | 25 | 74 | -196 |
| Broadleaf Deciduous Forest | 1994 | 627 | 637 | -2 | 619 | 548 | 11 |
| Broadleaf Evergreen Forest | 6921 | 1595 | 735 | 54 | 2024 | 1433 | 29 |
| Needleleaf Forest | 4599 | 1359 | 1006 | 26 | 1395 | 1160 | 17 |
| Mixed Forest | 966 | 310 | 246 | 21 | 311 | 285 | 8 |
| Mosaic Forest | 934 | 207 | 160 | 23 | 229 | 194 | 15 |
| Flooded Forest | 540 | 148 | 97 | 34 | 162 | 136 | 16 |
| Shrubland | 4226 | 912 | 643 | 29 | 1135 | 864 | 24 |
| Wetland | 468 | 144 | 74 | 49 | 159 | 107 | 33 |
| Barren | 1053 | 207 | 79 | 62 | 208 | 109 | 48 |
| **Total** | **31615** | **7055** | **5174** | **27** | **8025** | **6586** | **18** |

**Figure 3.**
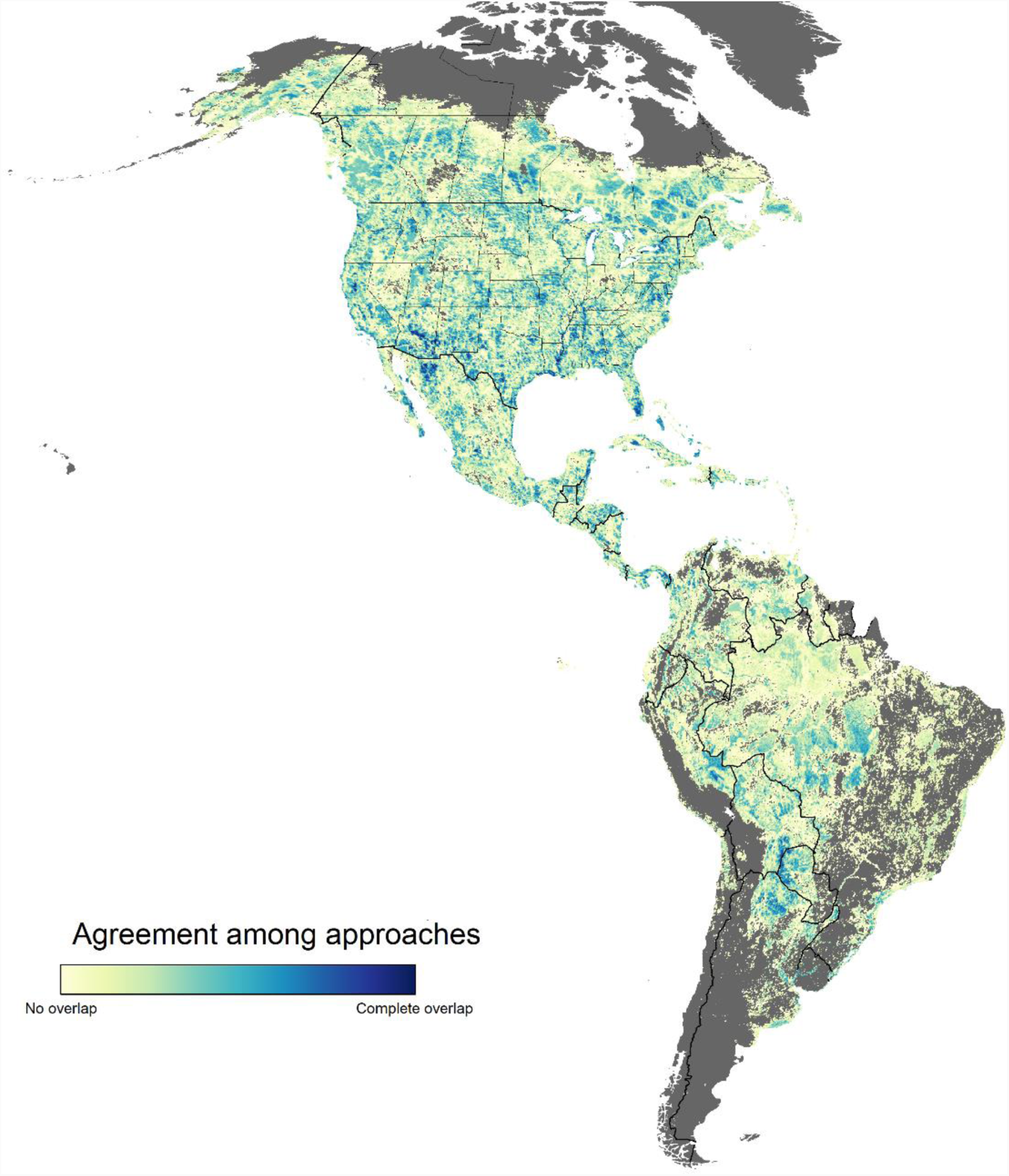
Range of agreement between the eight scenarios investigated. Darker blue indicates that most or all scenarios selected specific areas across the Western Hemisphere, and lighter yellow indicates areas of high scenario specificity. Scenario types considered: i) summing scenarios for each species in each week of the year vs. optimizing over all weeks and species in a full annual cycle, ii) including vs. ignoring spatial variation in population structure and migratory connectivity, and iii) incorporating vs. avoiding human-dominated landscapes in solutions.

Several additional caveats arise from our results, particularly with respect to land-sharing and sparing. Implementing conservation action in working landscapes may be more challenging than in areas with less human activity if the opportunity costs of management are higher in developed than undeveloped landscapes. For example, even if identified as a high-priority site for conservation in our land-sharing scenarios, land already converted to human use may be more vulnerable to degradation in the future than more intact areas^26^. Such habitat degradation, especially if combined with other anthropogenic stressors that may directly or indirectly reduce survival or performance of wildlife^27^, could make it difficult to reach population goals for species even if area needs are lower compared to less developed landscapes. In practice, both approaches are likely to be utilized given that target species will differ in their reliance on more or less developed habitats^28^. Therefore, our approach to prioritization provides planners with guidance on the approximate locations and requirements for land needed to meet our stated targets under a range of scenarios. With such portfolios in hand, planners can then more readily assess the cost-effectiveness of alternate approaches to land management and socio-economic policies most favorable to conservation and human well-being^21–23^. We also emphasize that the 30% target used here is illustrative only. In some cases, higher targets may be needed to avoid range contraction or the local extinction of sub-populations, to conserve ecological function such as seed dispersal or pest control^29^, or to maintain the evolutionary potential of locally-adapted populations^25,30^. Nevertheless, our 30% target returned solutions in all cases which vastly exceed the areal extent of existing conservation plans in support of Neotropical migrant birds.

## Conclusion

Ongoing declines in the abundance and distribution of many migratory species amid severe constraints on financial and human resources^31^ points to an urgent need for area-based plans that optimize the efficiency of conservation investments in ways that achieve conservation targets while minimizing the opportunity costs of land conservation and impacts on human livelihood^21–23,32^. Our solutions minimized the total land area prioritized for conservation to provide an area-efficient portfolio of lands for further consideration by conservation planners. Three key lessons can be derived from our results. First, scenarios based on the distributions of abundance of all 117 species over the entire annual cycle required less land area to meet conservation targets than scenarios based on optimizations that used the weekly distributions of those species throughout the year. Second, accounting for population structure through stratified sampling across the entire distribution of species increased the total land area required to achieve conservation targets. Despite requiring more land area, ensuring geographic representation may be necessary to the long-term persistence of species, particularly in widely-distributed species with population genetic structure potentially reflecting local adaptation to climatic conditions^25,30^. Third, area-based plans that accommodated human activity (land-sharing) were more efficient than land-sparing approaches that avoided areas with a high human footprint. However, because migrants vary spatially and temporally in their tolerance of human-impacted landscapes^33^, achieving conservation goals will likely require a portfolio of sites located in both intact and disturbed landscapes. Third, although our planning scenarios focused on Neotropical migratory birds, our approach could be easily adjusted and replicated in other migratory species and systems with sufficient data. In the case of birds, citizen science data and advanced prioritization tools allowed us to reveal marked efficiencies in area-based plans spanning the full annual cycle and multiple jurisdictions to conserve 117 individual species simultaneously.

## Methods

### Species selection

We included 117 species of Neotropical migratory passerines for our analysis (Supplemental Information Table 1). These species fell into two broad groups based on their breeding and stationary non-breeding ranges: 1) species where individuals breed in North America north of the US-Mexico border and migrate south of the Tropic of Cancer during the non-breeding period (n=101 species, SI Table 1), and 2) species with both migratory and resident populations or subspecies, for which individuals from migratory populations north of the US-Mexico border move south of the Tropic of Cancer during the non-breeding period (n=16 species).

### Approaches to conservation prioritization

We created 8 planning scenarios using weekly STEM models for each of 117 focal species and incorporating different assumptions about temporal scale and cost metrics employed in prioritization. First, we contrasted scenarios optimizing during each week of the year separately versus simultaneously over the entire annual cycle. We next created area-optimized solutions to conserve 30% of the global populations of all species in each week by sampling each species a) over their entire range, without accounting for population structure, or b) as 5 regional population clusters identified weekly to accommodate spatial variation in population structure and migratory connectivity. Third, we compared area-based conservation plans designed to represent different perspectives about the potential contribution of human-modified landscapes to the conservation of migratory birds, while including either the unrestricted cost metric or the human footprint cost metric, to create a total of 8 scenarios (SI Fig. 3). We used the prioritzr^34^ R package for the analysis, which interfaces with the Gurobi^35^ optimization software.

### Spatial prioritization approach

Here we use the concept of systematic conservation planning^36^, to inform choices about areas to protect, in order to optimize outcomes for biodiversity while minimizing societal costs^37^. To achieve the goal to optimize the trade-off between conservation benefit and socioeconomic cost, i.e. to get the most benefit for limited conservation funds, we strive to minimize an objective function over a set of decision variables, subject to a series of constraints. Integer linear programming (ILP) is the subset of optimization algorithms used here to solve reserve design problems. The general form of an ILP problem can be expressed in matrix notation as:

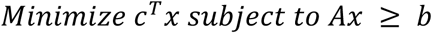

Where x is a vector of decision variables, c and b are vectors of known coefficients, and A is the constraint matrix. The final term specifies a series of structural constraints where relational operators for the constraint can be either ≥ the coefficients. In the minimum set cover problem, c is a vector of costs for each planning unit, b a vector of targets for each conservation feature, the relational operator would be ≥ for all features, and A is the representation matrix with Aij=rij, the representation level of feature i in planning unit j. We set an objective to find the solution that fulfills all the targets and constraints for the smallest area, which we use as our measure of cost ^11^. This objective is similar to that used in Marxan, the most widely used spatial conservation planning tool^38^.

### Spatiotemporal exploratory models

We used spatiotemporal exploratory models (STEM)^9,13,39^ to generate estimates of relative abundance for each species. STEM is a type of species distribution model created as an ensemble of local regression models generated from a spatiotemporal block subsampling design. Repeatedly subsampling and partitioning the study extent into grids of spatiotemporal blocks, and then fitting independent regression models (base models) in each block produces an ensemble of partially overlapping local models. Estimates at a given location and date are made by averaging across all the local models that contain the location and date. Combining estimates across the ensemble controls for inter-model variability^40^ and adapts to non-stationary predictor– response relationships^13^. To account for spatial variation in the density of the bird observation data^41^, smaller spatiotemporal blocks (10° × 10° × 30 continuous days) were used north of 12° latitude and larger blocks (20° × 20° × 30 continuous days) were used in the southern portion of the study extent.

The bird observation data used to implement STEM came from the eBird citizen-science database^42^. The data included species counts from complete checklists collected under the “traveling”, “stationary”, and “areal” protocols from January 1, 2004 to December 31, 2016 within the spatial extent bounded by 180° to 30° W Longitude (as well as Alaska between 150° E and 180° E). This resulted in a dataset consisting of 14 million checklists collected at 1.7 million unique locations, of which 10% were withheld for model validation.

Within each base model, species’ occupancy and abundance was assumed to be stationary. We fit zero-inflated boosted regression trees^9^ to predict the observed counts (abundance) of species based on three general classes of predictors: i) spatial predictors to account for spatial (and spatiotemporal) patterns; ii) temporal predictors to account for trends; and iii) predictors that describe the observation/detection process, which account for variation in detection rates, a nuisance when making inference about species occupancy and abundance. Spatial information was captured using elevation^43^ and NASA MODIS land^44^ and water cover data. The MODIS data were summarized as the proportion and spatial configuration of each of the 19 cover classes within 2.8 × 2.8 km (784 hectare) pixels centered at each eBird location using FRAGSTATS^45^ and SDMTools^46^. Summarizing the land-cover information at this resolution reduced the impact of erroneous cover classifications, and reduced the impact of inaccurate eBird checklist locations. The time of day was used to model variation in availability for detection; e.g., diurnal variation in behavior, such as participation in the “dawn chorus”^47^. Day of the year (1-366) was used to capture day-to-day changes in occupancy, and year was included to account for year-to-year differences. Finally, to account for variation in detection rates variables for the number of hours spent searching for species, the length of the transect traveled during the search, and the number of people in the search party were included in each base model.

Estimates of relative abundance and occupancy were rendered at weekly temporal resolution and 8.4 × 8.4 km spatial resolution. Each estimate was calculated as an ensemble average across 50-100 base models. The quantity estimated was either the expected number of birds of a given species (abundance) or the probability of the species being reported (occupancy) by a typical eBird participant on a search starting from the center of the pixel from 7:00 to 8:00 AM while traveling 1 km.

### Sampling for Population Structure and Migratory Connectivity

Many of the species used here are represented by multiple sub-species or populations known or suspected to follow different migratory pathways and use different breeding or wintering habitats^5,18,48^. However, in the absence of detailed knowledge on migration pathways for the vast majority of species, we developed a system of stratified sampling to represent the weekly distribution and spatial structure of each of 117 focal species to insure representation across their range throughout the annual cycle. To do so, we first conducted cluster analyses of weekly distribution maps for all 117 species to identify 5 clusters of equal abundance that encompassed the entire species range to insure representation across it. Our cluster analysis was based on a dissimilarity matrix of geographic locations and abundances (which were weighted by 1/3 to primarily focus on geographic effects and not bias cluster delineation toward spatially separated abundance clusters), and used the CLARA algorithm, which is an extension of the k-medoids technique for large datasets^49^. Our use of 5 clusters was arbitrary but flexible, and could be adjusted by the number of sub-species, races or sub-populations of interest.

### Land use constraints

We used two metrics to constrain our systematic conservation prioritization. First, we used a constant cost metric, where each planning unit was assigned a cost value of 1. Second, we used human footprint (2009; 1 km resolution)^19^ to identify areas more and less subject to human use, access or development pressures; specifically, we calculated the mean human footprint value for each 8.4 x 8.4 km pixel in our study area and used it as the ‘cost’ of each pixel during prioritization.

### Land cover representation

After the prioritization analyses, we summarized the major land cover types for each scenario that we generated. We used the 2015 data set of the global land cover map^50^ at a 300m resolution and clipped the original data to the study area. For each scenario, we used the geospatial data abstraction library^51^ to warp the selected cells from the prioritization onto the raster grid of the land cover dataset. There were 37 land cover classes identified across scenarios and the frequency and area amount of each was summarized for all scenarios. As a final step we combined similar land cover classes into broader classes (SI Table 2) and we used these to examine differences in area and land cover types selected under single season vs. full annual cycle planning and for land sparing vs land sharing scenarios (Table 2).

### Code availability

All computer code used in analysis, files generated from the analysis and outputs such as figures and tables have been deposited and will be made publicly available on publication here: https://osf.io/58hgs/?view_only=4bddcf37b95e470da3d3d90ba0f260de. The STEM model outputs used as inputs to the analysis will be made publicly available shortly by the Cornell Lab of Ornithology.

## Supporting information

## Acknowledgments

RS is supported by a Liber Ero Fellowship, ADR by a Garvin endowment, and JRB and PA by the Natural Sciences and Engineering Research Council of Canada. We also thank the eBird participants for their contributions and eBird team for their support. This work was funded by The Leon Levy Foundation, The Wolf Creek Charitable Foundation, NASA (NNH12ZDA001N-ECOF), Microsoft Azure Research Award (CRM: 0518680), and the National Science Foundation (ABI sustaining: DBI-1356308; computing support from CNS-1059284 and CCF-1522054). Data, analysis scripts and full results are archived here: https://osf.io/58hgs/?view_only=4bddcf37b95e470da3d3d90ba0f260de.

## Author Contributions

RS, SW, ADR, JRB and PA conceived the study. RS, DF and TA collected data and conducted analyses. All authors contributed to writing and editing the paper.

