## Supplementary material for "Optimizing the conservation of migratory species over their full annual cycle"

**Supplemental Information Table 1**. The 117 species of Neotropical migratory passerines considered for analysis and their range category. See Methods for range category descriptions. Species with an asterisk after the common name were left out of the analysis because of insufficient data for the stationary non-breeding range.

| **Common name** | **Scientific name** | **Range category** |
|---|---|---|
| Acadian Flycatcher | *Empidonax virescens* | 1 |
| American Redstart | *Setophaga ruticilla* | 1 |
| Ash-throated Flycatcher | *Myiarchus cinerascens* | 1 |
| Baltimore Oriole | *Icterus galbula* | 1 |
| Bank Swallow | *Riparia riparia* | 1 |
| Barn Swallow | *Hirundo rustica* | 2 |
| Bay-breasted Warbler | *Setophaga castanea* | 1 |
| Bell's Vireo | *Vireo bellii* | 1 |
| Black Swift | *Cypseloides niger* | 2 |
| Black-and-white Warbler | *Mniotilta varia* | 1 |
| Black-billed Cuckoo | *Coccyzus erythropthalmus* | 1 |
| Blackburnian Warbler | *Setophaga fusca* | 1 |
| Black-capped Vireo | *Vireo atricapilla* | 1 |
| Black-chinned Hummingbird | *Archilochus alexandri* | 1 |
| Black-headed Grosbeak | *Pheucticus melanocephalus* | 2 |
| Blackpoll Warbler | *Setophaga striata* | 1 |
| Black-throated Blue Warbler | *Setophaga caerulescens* | 1 |
| Black-throated Gray Warbler | *Setophaga nigrescens* | 1 |
| Black-throated Green Warbler | *Setophaga virens* | 1 |
| Blue Grosbeak | *Passerina caerulea* | 2 |
| Blue-gray Gnatcatcher | *Polioptila caerulea* | 2 |
| Blue-headed Vireo | *Vireo solitarius* | 1 |
| Blue-winged Warbler | *Vermivora cyanoptera* | 1 |
| Bobolink | Dolichonyx oryzivorus | 1 |
| Broad-tailed Hummingbird | *Selasphorus platycercus* | 1 |
| Bullock's Oriole | *Icterus bullockii* | 1 |
| Calliope Hummingbird | *Selasphorus calliope* | 1 |
| Canada Warbler | *Cardellina canadensis* | 1 |
| Cape May Warbler | *Setophaga tigrina* | 1 |
| Cassin's Kingbird | *Tyrannus vociferans* | 2 |

| | | |
|---|---|---|
| Cassin's Vireo | *Vireo cassinii* | 1 |
| Cave Swallow | *Petrochelidon fulva* | 2 |
| Cerulean Warbler | *Setophaga cerulea* | 1 |
| Chestnut-sided Warbler | *Setophaga pensylvanica* | 1 |
| Chimney Swift | *Chaetura pelagica* | 1 |
| Cliff Swallow | *Petrochelidon pyrrhonota* | 1 |
| Common Nighthawk | *Chordeiles minor* | 1 |
| Common Yellowthroat | *Geothlypis trichas* | 1 |
| Connecticut Warbler | *Oporornis agilis* | 1 |
| Cordilleran Flycatcher | *Empidonax occidentalis* | 1 |
| Dickcissel | *Spiza americana* | 1 |
| Dusky Flycatcher | *Empidonax oberholseri* | 1 |
| Eastern Kingbird | *Tyrannus tyrannus* | 1 |
| Eastern Wood-Pewee | *Contopus virens* | 1 |
| Golden-cheeked Warbler | *Setophaga chrysoparia* | 1 |
| Golden-winged Warbler | *Vermivora chrysoptera* | 1 |
| Grace's Warbler | *Setophaga graciae* | 2 |
| Gray Catbird | *Dumetella carolinensis* | 1 |
| Gray Vireo | *Vireo vicinior* | 1 |
| Gray-cheeked Thrush | *Catharus minimus* | 1 |
| Great Crested Flycatcher | *Myiarchus crinitus* | 1 |
| Hammond's Flycatcher | *Empidonax hammondii* | 1 |
| Hepatic Tanager | *Piranga flava* | 2 |
| Hermit Warbler | *Setophaga occidentalis* | 1 |
| Hooded Oriole | *Icterus cucullatus* | 2 |
| Hooded Warbler | *Setophaga citrina* | 1 |
| Indigo Bunting | *Passerina cyanea* | 1 |
| Kentucky Warbler | *Geothlypis formosa* | 1 |
| Lazuli Bunting | *Passerina amoena* | 1 |
| Least Flycatcher | *Empidonax minimus* | 1 |
| Louisiana Waterthrush | *Parkesia motacilla* | 1 |
| Lucy's Warbler | *Oreothlypis luciae* | 1 |
| MacGillivray's Warbler | *Geothlypis tolmiei* | 1 |
| Magnolia Warbler | *Setophaga magnolia* | 1 |
| Mourning Warbler | *Geothlypis philadelphia* | 1 |
| Nashville Warbler | *Oreothlypis ruficapilla* | 1 |
| Northern Parula | *Setophaga americana* | 1 |
| Northern Rough-winged Swallow | *Stelgidopteryx serripennis* | 1 |

| | | |
|---|---|---|
| Northern Waterthrush | *Parkesia noveboracensis* | 1 |
| Olive-sided Flycatcher | *Contopus cooperi* | 1 |
| Orange-crowned Warbler | *Oreothlypis celata* | 1 |
| Orchard Oriole | *Icterus spurius* | 1 |
| Ovenbird | *Seiurus aurocapilla* | 1 |
| Pacific-slope Flycatcher | *Empidonax difficilis* | 1 |
| Painted Bunting | *Passerina ciris* | 1 |
| Painted Redstart | *Myioborus pictus* | 2 |
| Palm Warbler | *Setophaga palmarum* | 1 |
| Philadelphia Vireo | *Vireo philadelphicus* | 1 |
| Plumbeous Vireo | *Vireo plumbeus* | 1 |
| Prairie Warbler | *Setophaga discolor* | 1 |
| Prothonotary Warbler | *Protonotaria citrea* | 1 |
| Purple Martin | *Progne subis* | 1 |
| Red-eyed Vireo | *Vireo olivaceus* | 2 |
| Red-faced Warbler | *Cardellina rubrifrons* | 2 |
| Rose-breasted Grosbeak | *Pheucticus ludovicianus* | 1 |
| Ruby-crowned Kinglet | *Regulus calendula* | 1 |
| Ruby-throated Hummingbird | *Archilochus colubris* | 1 |
| Rufous Hummingbird | *Selasphorus rufus* | 1 |
| Scarlet Tanager | *Piranga olivacea* | 1 |
| Scissor-tailed Flycatcher | *Tyrannus forficatus* | 1 |
| Scott's Oriole | *Icterus parisorum* | 2 |
| Summer Tanager | *Piranga rubra* | 1 |
| Swainson's Thrush | *Catharus ustulatus* | 1 |
| Swainson's Warbler | *Limnothlypis swainsonii* | 1 |
| Tennessee Warbler | *Oreothlypis peregrina* | 1 |
| Townsend's Warbler | *Setophaga townsendi* | 1 |
| Tree Swallow | *Tachycineta bicolor* | 1 |
| Vaux's Swift | *Chaetura vauxi* | 2 |
| Veery | *Catharus fuscescens* | 1 |
| Violet-green Swallow | *Tachycineta thalassina* | 1 |
| Virginia's Warbler | *Oreothlypis virginiae* | 1 |
| Warbling Vireo | *Vireo gilvus* | 1 |
| Western Kingbird | *Tyrannus verticalis* | 1 |
| Western Tanager | *Piranga ludoviciana* | 1 |
| Western Wood-Pewee | *Contopus sordidulus* | 1 |
| White-eyed Vireo | *Vireo griseus* | 1 |

| | | |
|---|---|---|
| White-throated Swift | *Aeronautes saxatalis* | 2 |
| Wilson's Warbler | *Cardellina pusilla* | 1 |
| Wood Thrush | *Hylocichla mustelina* | 1 |
| Worm-eating Warbler | *Helmitheros vermivorum* | 1 |
| Yellow Warbler | *Setophaga petechia* | 2 |
| Yellow-bellied Flycatcher | *Empidonax flaviventris* | 1 |
| Yellow-billed Cuckoo | *Coccyzus americanus* | 1 |
| Yellow-breasted Chat | *Icteria virens* | 1 |
| Yellow-rumped Warbler | *Setophaga coronata* | 1 |
| Yellow-throated Vireo | *Vireo flavifrons* | 1 |
| Yellow-throated Warbler | *Setophaga dominica* | 1 |

6

**Supplemental Information Table 2**. Summarized and individual land cover classes used to examine selection of land cover types under single season vs. full annual cycle planning and for land sharing vs sparing scenarios. See ref. 44 for further descriptions of the individual land cover classes.

| **Summary class** | **Individual classes** |
|---|---|
| Cropland/Mosaic Cropland | Cropland/rainfed |
| Cropland/Mosaic Cropland | Cropland, irrigated or post-flooding |
| Cropland/Mosaic Cropland | Mosaic cropland (>50%) / natural vegetation (tree, shrub, herbaceous cover) (<50%) |
| Cropland/Mosaic Cropland | Mosaic natural vegetation (tree, shrub, herbaceous cover) (>50%) / cropland (<50%) |
| Grassland | Herbaceous cover |
| Grassland | Grassland |
| Urban areas | Urban areas |
| Broadleaf Deciduous Forest | Tree cover, broadleaved, deciduous, closed to open (>15%) |
| Broadleaf Deciduous Forest | Tree cover, broadleaved, deciduous, closed (>40%) (61) |
| Broadleaf Deciduous Forest | Tree cover, broadleaved, deciduous, open (15-40%) (62) |
| Broadleaf Evergreen Forest | Tree cover, broadleaved, evergreen, closed to open (>15%) |
| Needleleaf Forest | Tree cover, needleleaved, evergreen, closed to open (>15%) |
| Needleleaf Forest | Tree cover, needleleaved, evergreen, closed (>40%) (71) |
| Needleleaf Forest | Tree cover, needleleaved, evergreen, open (15-40%) (72) |
| Needleleaf Forest | Tree cover, needleleaved, deciduous, closed to open (>15%) |
| Needleleaf Forest | Tree cover, needleleaved, deciduous, closed (>40%) (81) |
| Needleleaf Forest | Tree cover, needleleaved, deciduous, open (15-40%) (82) |
| Mixed Forest | Tree cover, mixed leaf type (broadleaved and needleleaved) |
| Mosaic Forest | Mosaic tree and shrub (>50%) / herbaceous cover (<50%) |
| Mosaic Forest | Mosaic herbaceous cover (>50%) / tree and shrub (<50%) |
| Flooded Forest | Tree cover, flooded, fresh or brakish water |
| Flooded Forest | Tree cover, flooded, saline water |
| Shrubland | Shrubland |
| Shrubland | Shrubland evergreen |
| Shrubland | Shrubland deciduous |
| Wetland | Shrub or herbaceous cover, flooded, fresh/saline/brackish water |
| Barren | Lichens and mosses |
| Barren | Sparse vegetation (tree, shrub, herbaceous cover) (<15%) |
| Barren | Bare areas |
| Barren | Consolidated bare areas |
| Barren | Unconsolidated bare areas |

**Supplemental Information Table 3.** Area selected (1000 $km^2$) for major land cover types using weekly planning for land sharing vs. sparing scenarios and for single population vs population structure approaches. Area available is the total amount of each land cover available based on all cells throughout the year where ≥ 1 species was present. % reduction is the percentage decrease in the area required for each land cover type with full annual cycle compared to single season planning. Not all land cover classes are included in the table and therefore individual land cover values do not sum to the total in each column. Land cover data was extracted from the global land cover map for 2015 (300m resolution) [50].

| **Land cover** | **Area available** | **Single Population** | | | **Population Structure** | | |
|---|---|---|---|---|---|---|---|
| | | Land Sparing | Land Sharing | % reduction | Land Sparing | Land Sharing | % reduction |
| Cropland/Mosaic Cropland | 2269 | 596 | 895 | -50 | 875 | 1295 | -48 |
| Grassland | 5555 | 2091 | 3139 | -50 | 2387 | 3769 | -58 |
| Urban areas | 205 | 14 | 171 | -1121 | 53 | 186 | -250 |
| Broadleaf Deciduous Forest | 1994 | 1032 | 1385 | -34 | 1217 | 1611 | -32 |
| Broadleaf Evergreen Forest | 6921 | 3436 | 1606 | 53 | 3984 | 4064 | -2 |
| Needleleaf Forest | 4599 | 2882 | 2806 | 3 | 2792 | 3266 | -17 |
| Mixed Forest | 966 | 610 | 749 | -23 | 611 | 850 | -39 |
| Mosaic Forest | 934 | 381 | 368 | 3 | 431 | 527 | -22 |
| Flooded Forest | 540 | 287 | 230 | 20 | 312 | 369 | -18 |
| Shrubland | 4226 | 1763 | 1925 | -9 | 2169 | 2498 | -15 |
| Wetland | 468 | 219 | 161 | 26 | 287 | 276 | 4 |
| Barren | 1053 | 415 | 222 | 47 | 441 | 303 | 31 |
| **Total** | **31615** | **13727** | **13656** | **1** | **15558** | **19015** | **-22** |

**Supplemental Information Fig 1.** Species diversity based on relative abundance spatial temporal models of roughly 117 migratory songbird species is shown for every week of the year. Animated GIF uploaded as separate file. Here week 1 as a placeholder.

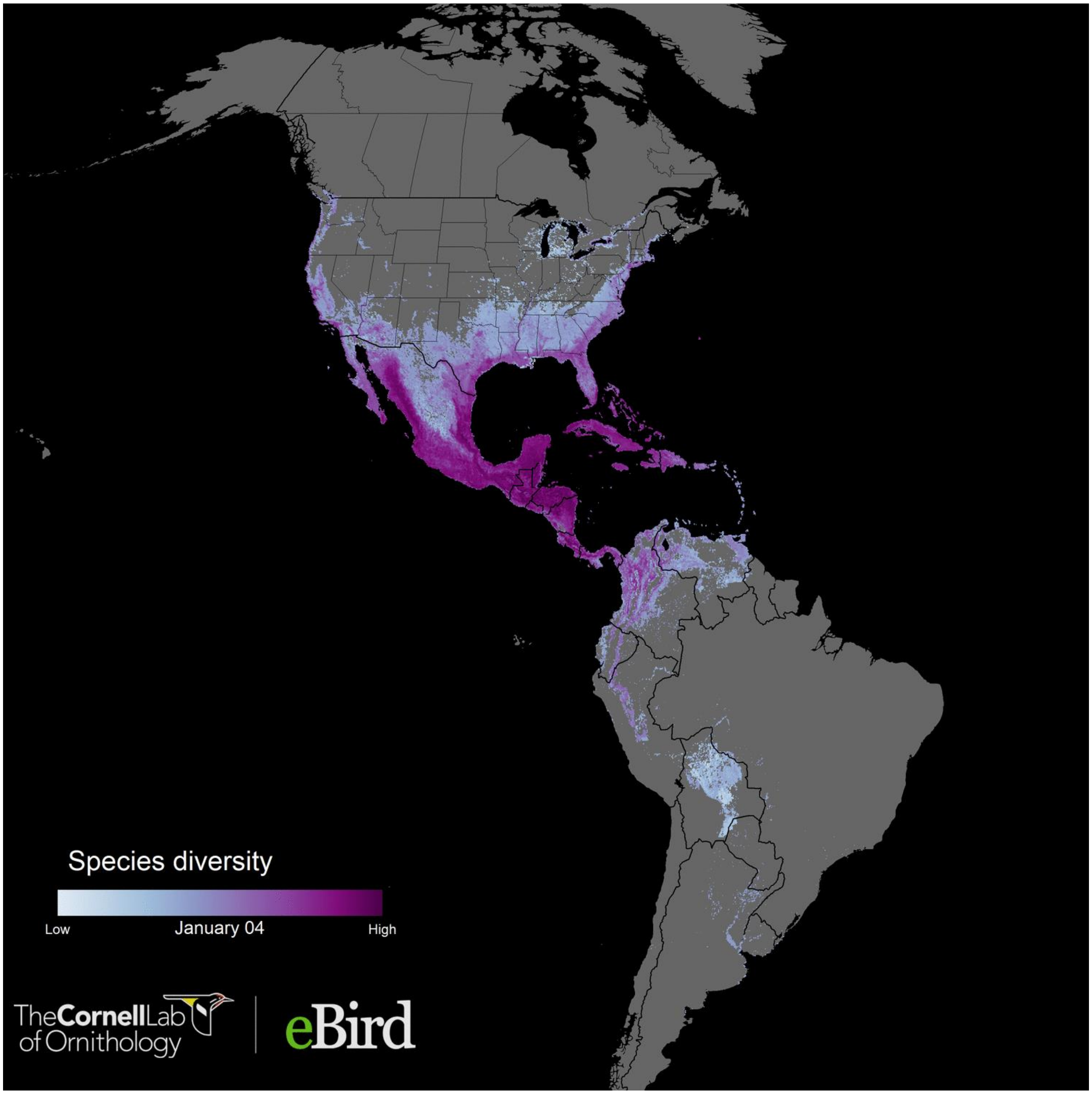

**Supplemental Information Fig 2**. Human footprint estimated in 2009[19].

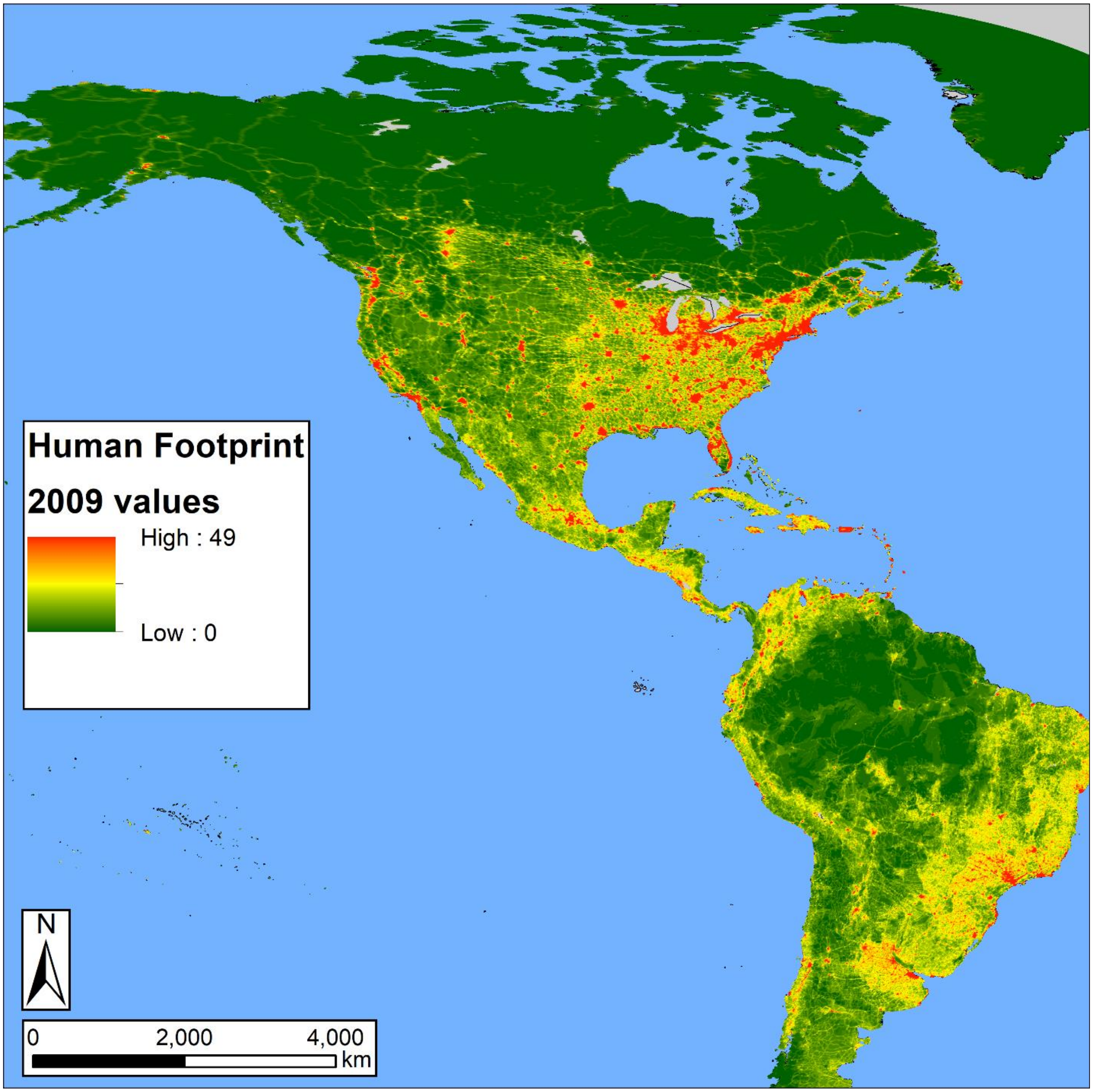

**Supplemental Information Fig 3.** Schematic of conservation prioritization scenarios investigated. Runs represents the number of prioritizations we completed per scenario. Boxes in green show the actual scenarios investigated in this study.

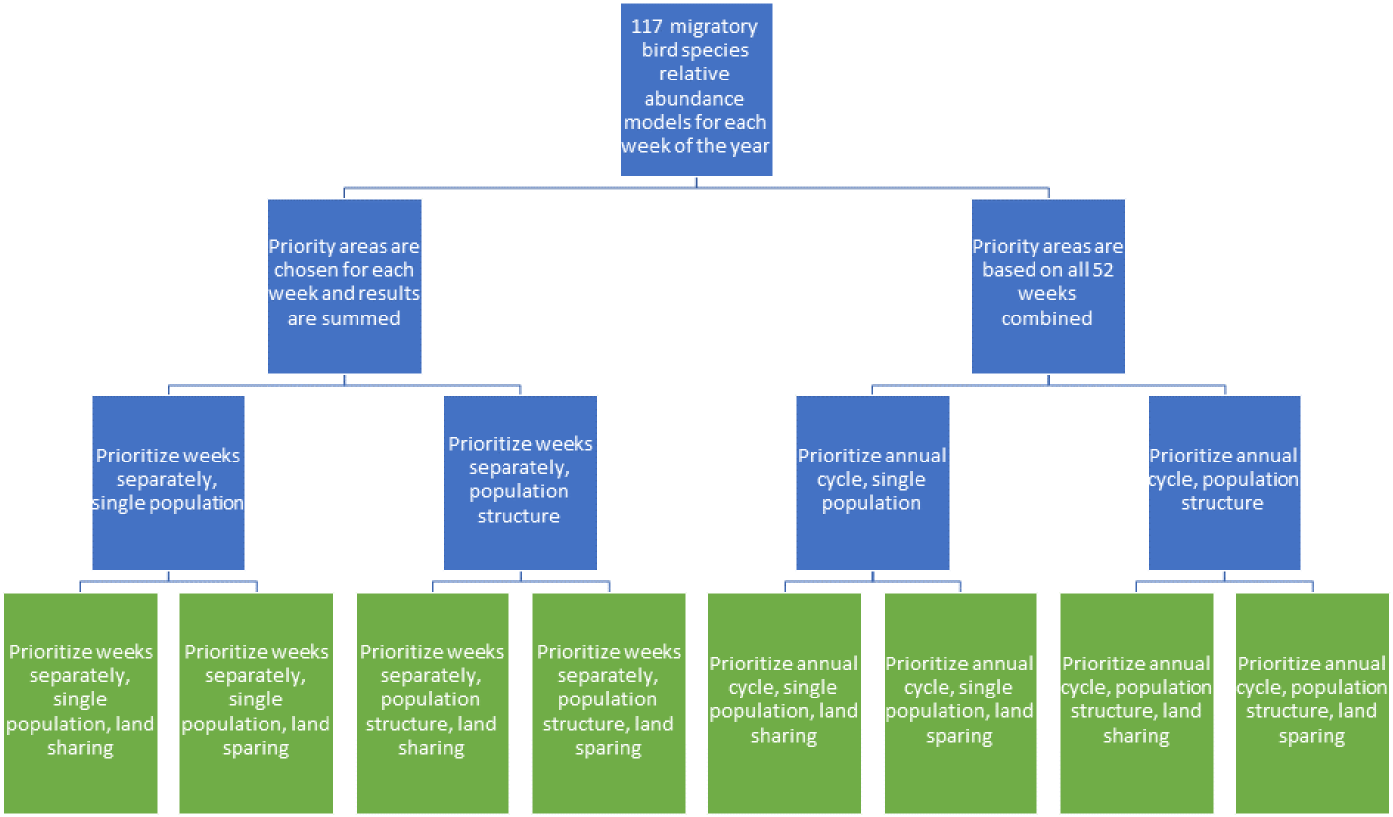

**Supplemental Information Fig 4.** Detailed version of Figure 1, focusing on northern South America. Comparison of areas prioritized for weekly and full annual cycle planning under a land sharing approach allowing for the inclusion of human dominated landscapes versus a land sparing approach that excludes areas of high human footprint. The prioritization is based on a target of 30% of global populations of 117 species of Neotropical migratory birds when each species range is considered as a single population. a) = land sharing, weekly, b) = sharing annual cycle, c) = land sparing weekly, d) = land sparing annual cycle.

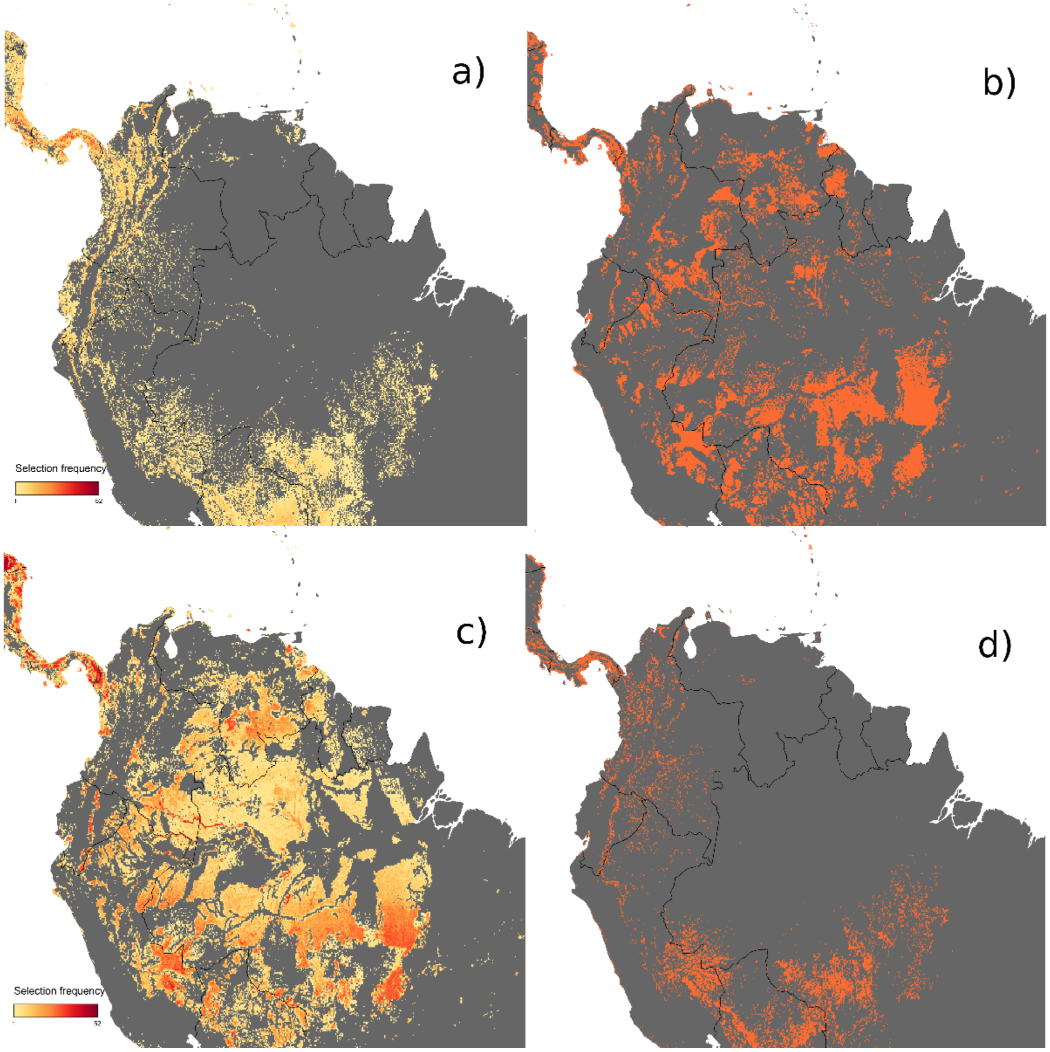


**Supplemental Information Fig 5.** Detailed version of Figure 2, focusing on northern South America. Comparison of areas prioritized for weekly and full annual cycle planning under a land sharing approach allowing for the inclusion of human dominated landscapes versus a land sparing approach that excludes areas of high human footprint. The prioritization is based on a target of 30% of global populations of 117 species of Neotropical migratory birds when each species range is considered with population structure (five regional clusters). a) = land sharing, weekly, b) = sharing annual cycle, c) = land sparing weekly, d) = land sparing annual cycle.

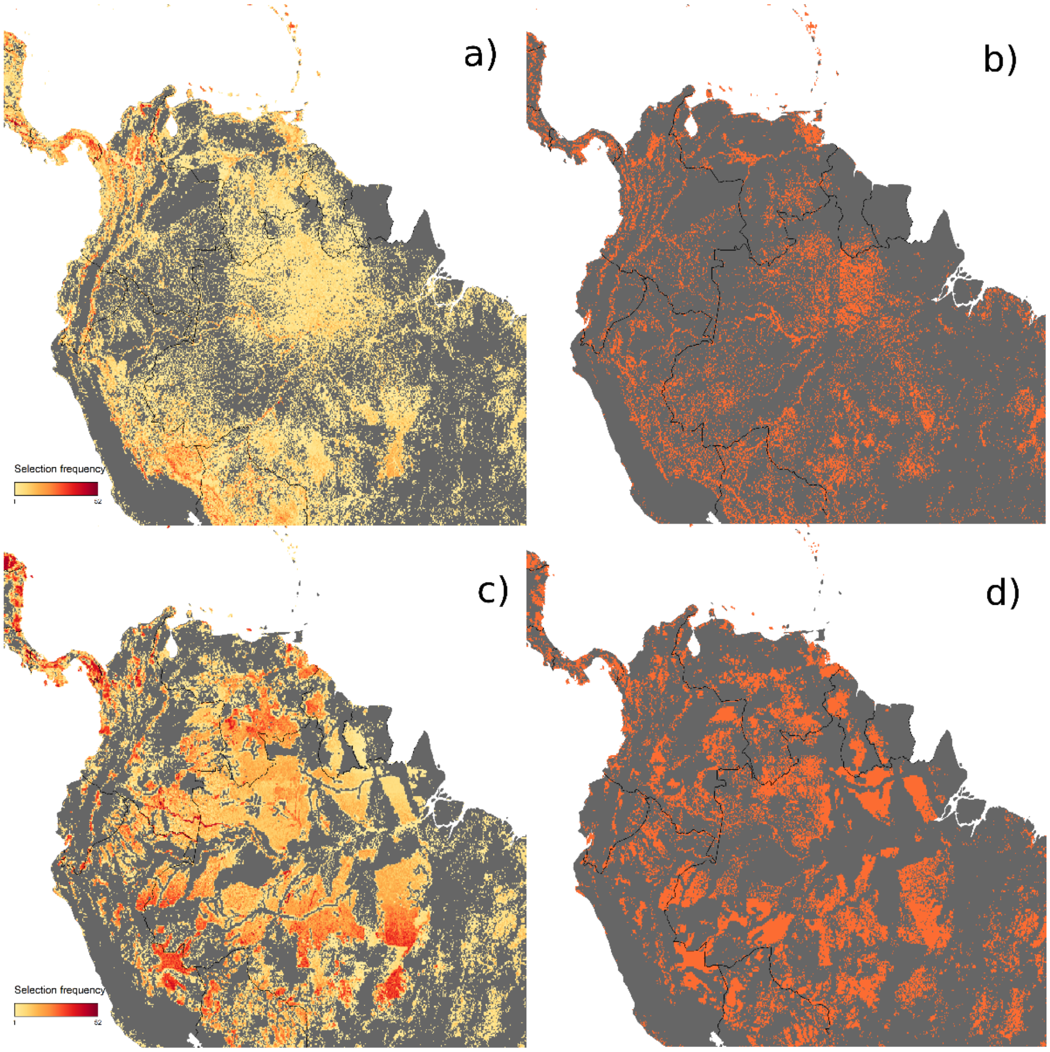
